# A new approach to design artificial 3D micro-niches with combined chemical, topographical and rheological cues

**DOI:** 10.1101/291104

**Authors:** Celine Stoecklin, Zhang Yue, Wilhelm W. Chen, Richard de Mets, Eileen Fong, Vincent Studer, Virgile Viasnoff

## Abstract

The *in vitro* methods to recapitulate environmental cues around cells are usually optimized to test a specific property of the environment (biochemical nature or the stiffness of the extra cellular matrix (ECM), or nanotopography) for its capability to induce defined cell behaviors (lineage commitment, migration). Approaches that combine different environmental cues in 3D to assess the biological response of cells to the spatial organization of different biophysical and biochemical cues are growingly being developed. We demonstrate how the lamination of through-hole polymeric bio-functionalized membranes can be implemented to create complex *bona fide* micro-niches with differential 3D environmental properties using photoactive materials. Our approach enables to create micro-niches ranging in size from single cells to cell aggregates. They are bio-functionalized in 3D simultaneously with topographical featured, protein patterns and structured ECM surrogate with 1 micrometer resolution. We demonstrate how these niches extend in 3D the ability to pattern cells. We exemplify how they can be used to standardize cells shapes in 3D and to trigger the apico-basal polarization of single epithelial cells.

## 1 Introduction

Our understanding of how cells sense the biochemical and biophysical properties of their environment is rapidly advancing. The nature and spatial structure of the extracellular matrix, the presence of cell-cell interactions, and the variation of confinement have emerged as important signaling factors that cells integrate to commit to a defined state.^[1-3]^ It results that large effort have been devoted to create screening methods to optimized the biochemical nature of the extracellular matrix (ECM), its stiffness or the topographical cues that favor the differentiation of stem cells into a given lineage. ^[4-7]^. *In vitro* approaches to create artificial micro-niche presenting combinations of environmental cues in 3D also advance our understanding of the molecular pathways involved in environmental sensing. ^[8,9]^ In this perspective, 2D protein patterning techniques provide a handy tool to promote cell adhesion on substrates of various rigidities and topographies.^[10-12]^ However extension of protein patterning to a 3D space is still limited.^[13,14]^ Instead, bio-functionalized hydrogels^[15],[16]^ or extracellular matrix (ECM) extracts ^[17]^ are tools of choice for “3D biology”, especially in the development of organoids.^[18-20]^. Bio-printing of ECM allows the spatial structuration of cell co-cultures in hydrogels at the hundred of micron scale.^[21]^ Two photons micro-fabrication provides sub-micron resolution to the expense of very slow output^[22]^. Microfluidic approaches have also been largely used to control flow and develop organ-on-chip devices.^[23,24]^. Many of these experimental approaches are dedicated to a specific environment type. We present here a unifying and versatile experimental approach to design 3D local microenvironments combining together 3D protein patterns, topographical features and structured hydrogels.

It relies on perforated bio-functionalized polymeric layers used as ‘Lego-like bricks’. After we detailed the fabrication steps, we provide the extensive list of cues that can be controlled. Then, we showcase a particular example where the niches can be used to pattern cells in 3D and induce the early stages of apico-basal polarization in hepatocytes by a proper combination of environmental cues.

## 2 Results

### 2.1 General fabrication and assembly principles

The general principle of our methods relies on the layer-by layer lamination of micro-fabricated membranes (**Figure 1a-d**). The layering process is key in creating micro-compartments with layered physical properties that can be bio-functionalized differently by protein adsorption, UV protein printing or localized hydrogel grafting. The membrane assembly is deposited in a Petri dish or in a flow chamber to enable cell culture (**Figure 1a-b**). **Figure 1c-d** shows the general principle to assemble a cavity comprising a perforated membrane (creating the pit wall) assembled onto a textured layer constituting the base and covered with a porous lid. We fabricated the perforated membranes (pit layers) in photo-curable polymers (NOA 73, or BIO-134). We used standard soft lithography techniques to produce silicon wafers with pits of defined shapes. To form the pit layer, the UV-curable commercial pre-polymer mix was infiltrated by capillarity in the interstitial space separating the negative PDMS replica and a flat PDMS substrate. We tuned UV exposure doses to create a self-standing membrane solid enough to be manipulated but still retaining surface chemical activity (**Table S2**). The unreacted acrylate moieties on surface are then readily available for subsequent grafting. We found the residual chemical activity higher for NOA 73 polymer than for the fluoro-containing polymer BIO-134. However, BIO-134 has an optical index matching cells and tissues (n=1.34). It is thus particularly favorable for fabricating nano/micro-structures and ensures superior imaging qualities of the micro-niches. Partial curing is also essential to ensure sufficient adhesiveness of the pit layer onto any other substrates without further chemical treatment. For the base layer, we tested rigid substrates (glass, plastic cell culture dishes), softer substrates (soft PDMS, hydrogels), or nano/micro-textured substrates (Nanosurface Biomedical, PDMS, PP, PET). Besides providing different chemical surface properties or topographies, these materials adequately bound to the pit layers. After cell seeding, the assembled membranes can either be immersed in cell culture medium, or covered with an ECM gel (Matrigel or collagen), which creates a porous lid on top of the wells.

**Figure 1.**
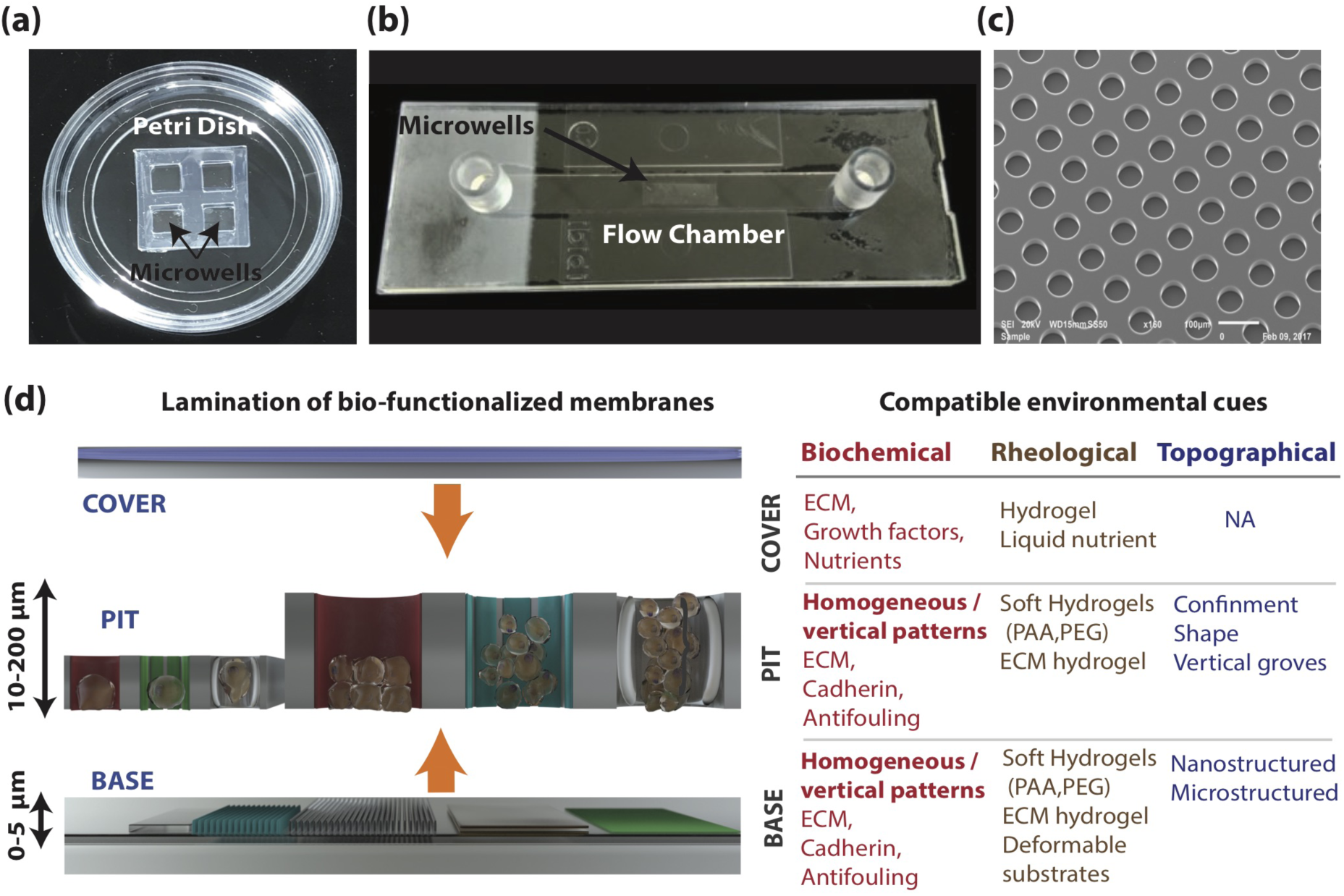
‘Lego-like’ approach to control micro-environmental cues in 3D. **(a)** Final assembly of the cell culture system in a petri dish with 4 chambers of microwells. **(b)** Alternative assembly of the microwells membrane in a flow chamber. **(c)** SEM images showing microwells of 60 µm in diameter and 20 µm in height made of NOA 73. **(d)** Left: General principle of assembly of the base, pit and cover layers, tuneable independently, that lead to the creation of a biocompatible micro-niches with different scales. Right: List of compatible biochemical, rheological, and topographical cues that can be combined among the three different layers.

### 2.2 3D Bio-functionalization with multiple protein patterns

The layered membranes provide a 2D scaffold that extends in volume. Adapting 2D multi-protein printing methods to our systems allows to bio-functionalize the surfaces in 3D. The pit and base membranes can be independently and homogeneously bio-functionalized prior to their assembly. **Figure 2a** depicts the method to create niches with homogeneous but distinct adhesive cues on the pit and bottom of the niche. Each layer was separately incubated in a solution containing ECM proteins or E-cadherin protein. The main steps are summarized below (**Note S2**). The base layer is functionalized by simple protein adsorption, followed by a rinsing and drying step. The pit layer (as large as a few cm^2^) is functionalized on a protective sacrificial substrate that restricts protein adsorption to the top of the membrane and inside the holes. The sacrificial substrate is then removed, and the pit layer flipped onto the base surface, thereby exposing an unreacted surface on the top. Using a solution of 0.2%(w/v) of pluronic-acid the exposed surface is passivated to ensure cells are seeded only inside the cavities. Using a single base and a single pit layer leads to niches with distinct proteins on their sides and bottom (**Figure 2b**). Repeating the layering process with multiple pit layers enables to create multi-tier niches with a vertical stratification of biochemical cues (**Figure 2c**). The approach described above limits the 3D protein patterns to horizontal stripes. More complex patterns can be achieved using the commercial 2D protein printing system PRIMO (Alveole). This system relies on the local adsorption of proteins to surfaces using UV light-induced molecular adsorption mediated by a benzophenon (PLPP).^[25]^ Here, UV patterns are directly projected onto a horizontal surface through a microscope, using a DMD (Digital Micro-mirror Device). **Figure 2d** summarizes the main steps involved in the printing process (**Note S2** and **Table S3**). The bare membranes are first assembled and passivated with PLL-g-PEG (poly-L-lysine-*graft*-polyethylene glycol). A homemade plugin is used to digitally align the projected patterns with the microwells image for each field of view. A single field of view of 640 µm x 1024 µm for 10 X magnification, 320 µm x 512 µm for 20 X magnification and 160 µm x 256 µm for 40 X can be printed at a time. Moving the microscope stage and running the digital alignment repeatedly allow the easy printing on multiple fields of view. It leads to printing time of around 4 minutes/mm^2^ for a 10 X magnification (XY resolution 2.4 µm), 6.1 minutes/mm^2^ for a 20X objective (XY resolution 1.2µm) and 16.6 minutes/mm^2^ for a 40X objective (XY resolution 600nm) for a single protein. Multi-protein printing is achieved by repeating the printing process with another protein. **Figure 2e** depicts a checkerboard pattern (20×5 µm to 3×3 µm), printed on hemispheric pits. The local curvature of the pit (compensated by the design of the projected pattern) ensures a vertical resolution equal to the lateral one (**Movie S1**). In the case of pits with vertical walls, the lateral resolution is conserved but the vertical resolution is ill defined and largely depends on the objective characteristics, the divergence of the pattern illumination and the exposure time. In this case, the vertical patterning is limited by the height of the well. **Figure 2f** illustrates a micro-niche (BIO-134, 40µm cube) printed with a 10 µm diameter circle at the bottom (fibronectin rhodamin) and 30 µm wide vertical stripes (Fibrinogen-Alexa 488). The coating conditions, and exposure-time for various objective magnifications, are summarized in **Table S3**. The printing process is fast enough to pattern arrays of thousands of niches in a couple of hours (**Figure 3**, N=1040 wells, 1.5h printing time) for a single protein. The well occupancy (**Figure S1**) reaches between 65 to 85% (N=1040 wells) depending on cell types and patterns. **Figure 3** represents seeding of HFF (human foreskin fibroblasts) with a well occupancy of 80%.

**Figure 2.**
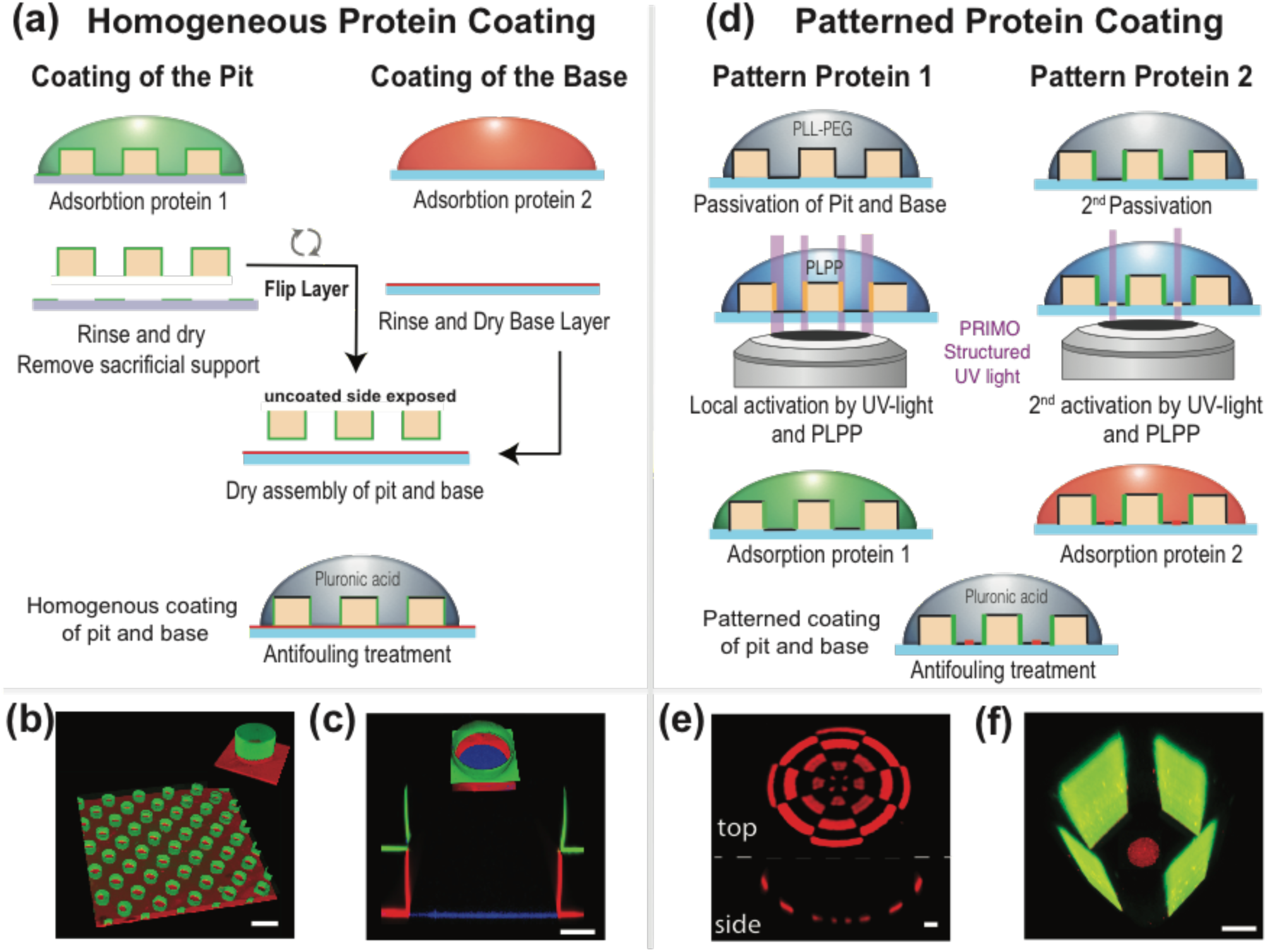
Microwells bio-functionalization with homogeneous and patterned protein coating techniques. **(a)** Schematic principle of differential and homogeneous protein coating on the pit and on the base layer. The pit and base are coated with different biochemical cues separately and then assembled and passivated to form the final micro-niche. **(b)** Example of differential homogeneous coating of 15µm MyPol-134 microwells with E-cadherin (green) on the pit and laminin rhodamine (red) on the base. (**c)** Cross-section and 3D visualisation of a 200 µm diameter micro-niche composed of two aligned pit layers of NOA 73 with differential homogeneous coating. Fibrinogen Alexa 488 (green) is coated on the top pit layer, Fibrinogen Alexa 546 (red) on the middle pit layer and Fibrinogen Alexa 647 (blue) on the base layer. **(d)** Schematic principle of patterned protein coating with PRIMO. Pit and base are first passivated with an antifouling poly-ethylene-glycol (PLL-g-PEG). Addition of PLPP photoinitiator allows under illumination with UV-light (375nm) to cleave the PEG chain at a chosen location. Incubation of the protein leads to a patterned coating of the micro-niche in 3D. The process can then be repeated to coat another protein of interest. **(e)** En face and side viewing of an hemispheric pit (50µm diameter and 30 µm height) of NOA 73 patterned with a fibronectin rhodamine checkerboard (red). **(f**) 3D visualisation of a MyPol-134 square micro-well (40 µm) coated with 30 µm stripes of Fibrinogen Alexa 488 (green) on the pit and a 10µm diameter circle of Fibronectin Rhodamine (red) on the base layer. Scale bars (**b**), (**c**), (**e**), (**f**): 20 µm.

**Figure 3.**
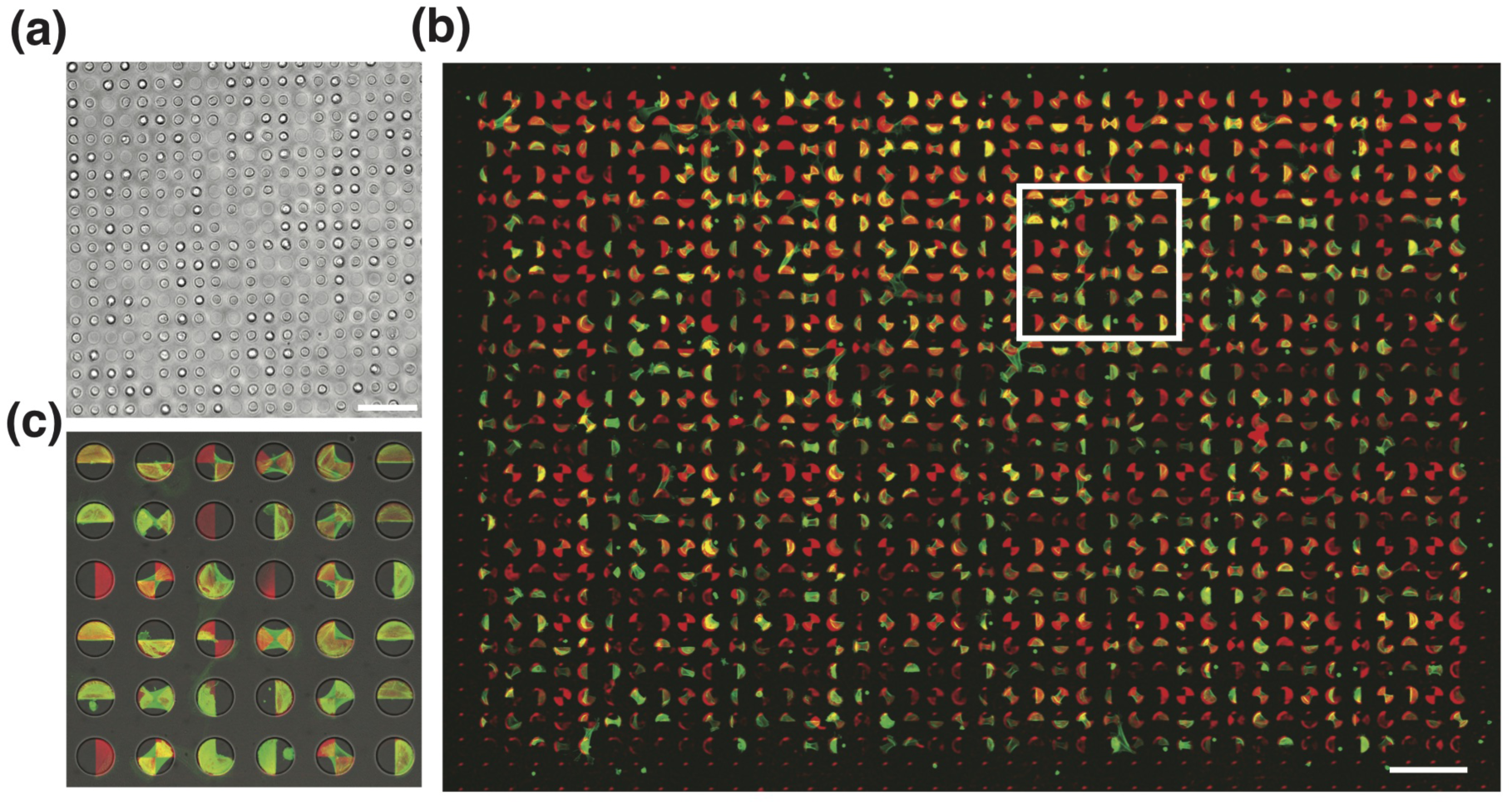
Single cells seeding and occupancy in the microwells. **(a)** Single cells seeding of Human Foreskin Fibroblast (HFF) in the microwells before attachment to the fibronectin patterned area. 80% of the microwells are filled with cells. (**b)** Confocal images of an array comprising of 1080 microwells (NOA 73, 60 µm diameter and 30 µm height) printed with Fibronectin Rhodamin patterns projected on the bottom and the wall of the microwells. HFF cells were cultured for one day on the patterns and stained for phalloidin-TRITC to visualize the cells shapes. The confocal images have been stitched together to reconstruct the whole chip array of 2430 µm x 3600 µm. **(c)** Zoom over the white square area of 510µm x 510µm. A phase contrast image has been overlayed to the confocal image for a better visualization of the microwells. From the 80% of the cells that attached to the patterns, 50% adopt the proper 3D patterning organization for analysis. Scale bar **(a), (b)**: 300 µm

### 2.3 Structuration of cell cytoskeleton, single cell apico-basal polarization

We then demonstrated that our technique enables the normalization of the cellular cortex in 3D. Classical 2D patterning allows the shape and size of the cells to be standardized by constraining their “adhesive imprint” on the substrate. Continuous 2D geometrical shapes (circle, squares) constrain the cell shape but usually do not normalize the spatial distribution of cytoskeletal structures (eg stress fibers). In contrast, discontinuous shapes (bow, arrow lines) define the structure of the adhesion area, and lead to the development of stereotypical fibers. ^[26,27]^ In 3D, the continuous patterning corresponds to the homogeneous coating of the well, which imposes its shape to the cell by physical ^[28]^ or adhesive interactions.^[29-32]^ Cytoskeletal organization however, is still irreproducible from one cell to the other. **Figure 4a** shows a 3D rendering of the average (N∼25) shape of primary rat hepatocytes in circular microwells patterned in 3D with fibronectin structured as a crossbow, a triangle and a cross (**Movie S2**). Full 3D stacks are presented in **Movie S3.** The cell displays reproducible actin accumulation zones, not only at the adhesive point but also over the suspended part of the cells. Vertical gradients of actin accumulation are also revealed along the suspended edges. These likely indicate the height dependent tension of the cortex. Note that the wells were fabricated in BIO-134 to avoid any deterioration of the image quality along the z axis, and to maintain maximal imaging quality of the cell/well interacting area. ^[33]^ **Figure 4b** also displays the distribution of nuclei localization relative to the average cell shape. Note that the hepatocytes nuclei locate few microns away from the pattern center (6.3 um ± 0.7µm for circle 3D patterns, 4.1 um ± 1µm for cross 3D patterns, 3.5 um ± 1.7µm for triangle 3D patterns and 4.4 um ± 1.6µm for crossbow 3D patterns). In the triangular and ‘cross’ pattern, the nuclei locate towards one vertex, whereas in the crossbow pattern the nuclei accommodate in the vicinity of the semi-circular area (crossbow head). Vertically, all hepatic nuclei are pushed down in contact with the substrate. We then compared the localization of the nuclei between hepatocytes and HaCaT (keratinocyte) cells. We modified the dimensions the wells to account for the difference in size of the hepatocytes and HaCaT cells in suspension. We found that cells would adopt the shaped guided by the patterns only if the dimensions of the microwell were within a specific range. If the wells are too small the cells can not enter, but if they are too big, the cells only adhere partially to the walls, and this is in a non-columnar manner. Compared to hepatocytes, the HaCaT cells possessed nuclei that were closer to the center of the pattern in the horizontal plane (**Figure 4c**). In the vertical direction hepatocyte nuclei lie close to the base of the microwell, whereas in HaCaT cells they were located in the vicinity of the median plane.

**Figure 4.**
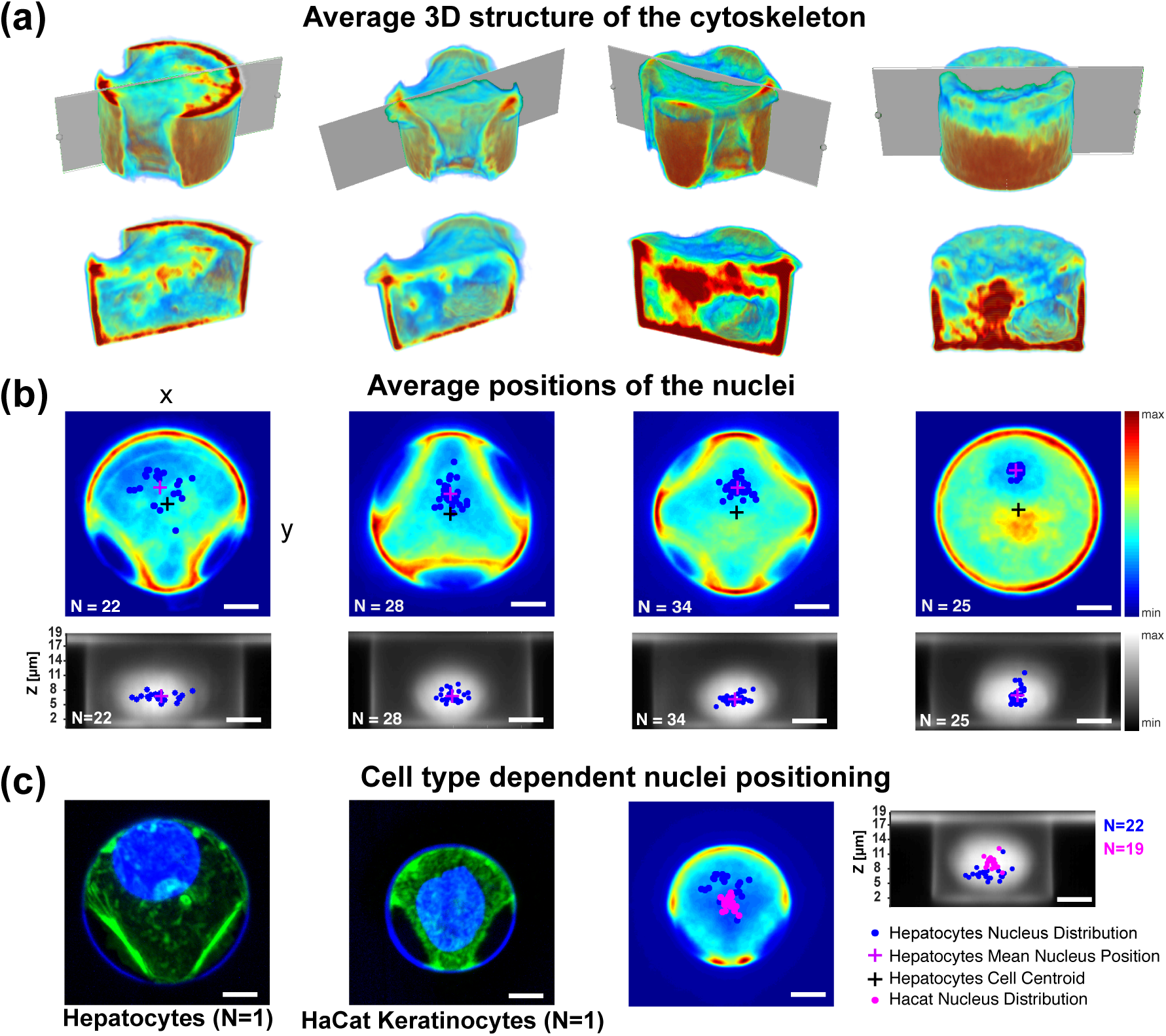
Structuration of the cell actin cytoskeleton in 3D on different geometrical shapes. (**a**) Average 3D actin structure of hepatocytes cytoskeleton on different fibronectin geometrical shapes (crossbow N=22, triangle N=28, cross N=34, circles N=25). The different geometrical shapes are patterned with PRIMO in 3D on 25µm diameter and 18µm deep microwells fabricated in BIO-134. **(b)** Comparison of the nucleus localisation in x,y,z between the different geometrical shapes for the hepatocytes. Top: Average actin intensity projected along the z axis superimposed with the nuclei distribution (blue dots) for each shape. Bottom: Average DAPI intensity projected along the y axis to visualize the nuclei height distribution in the micro-well. **(c)** Different cell types exhibit different nucleus positioning. Left: Maximum Actin (green) and DAPI (blue) intensity projection along z of one hepatocyte (N=1) and one HaCat cell (N=1). Right: Comparison between the HaCat nuclei distribution (magenta) and hepatocytes nuclei distribution (blue) in x,y and x,z. The hepatocytes nuclei distribution was normalized to account for the micro-well size difference between of the HaCat (20 µm diameter, 18µm deep) and hepatocytes (25 µm diameter, 18µm deep). The nuclei distribution in height for HaCat cells are significantly different than for the hepatocytes (p value <10^−3^). Scale bar (**b**), (**c**): 5 µm.

These results prompted us to consider our system as a good way to study the role of 3D biophysical cues in the development of apico-basal polarization (ABP) of epithelial cells. Nuclear positioning is a prominent hallmark of this ubiquitous biological process. To do this, we placed single primary rat hepatocytes in cavities coated with E-cadherin on their walls, and fibronectin on their base (E/F coating) as illustrated in **Figure 5a.** After fixation, we stained the cells to visualize the intracellular domains of E-cad (green), actin (red) and ZO1 (blue). 80% (N=117) of the cells (**Figure 5b**) displayed a discontinuous cadherin structure with an ellipsoidal region showing no cadherin recruitment (**Figure 5c, Movie S4**). A thick actin belt, which was associated with an accumulation of ZO1, systematically surrounds this ellipsoidal region. Radial projections on **Figure 5c** clearly demonstrate this stereotypic organization. Here, 7% of the cells exhibited zones lacking E-cad, and these were located opposite each other (**Figure 5c**). In 13% of the cases, cells had no specific structures and display a continuous cadherin distribution and homogeneous cortex with scattered puncta of ZO1. Ellipsoidal protein distribution was only visible in the E-F coated wells. Hepatocytes placed in microwells with homogeneous fibronectin coating (F/F coating) were 100% devoid of such ellipsoidal protein organization. This organization is reminiscent of intercellular lumen, or canaliculi, development between hepatocytes. In this case the canaliculi constitutes the apical pole of the cells. A single hepatocyte can form as many apical regions as it has neighbors. The complete biological description of how such single-cell-based structures are induced by the biophysical cues contained within the niche will be sought in future work. Taken as a whole, our data exemplifies the potential of our approach to guide and induce apico-basal polarization using defined external cues, at the single cell level.

**Figure 5.**
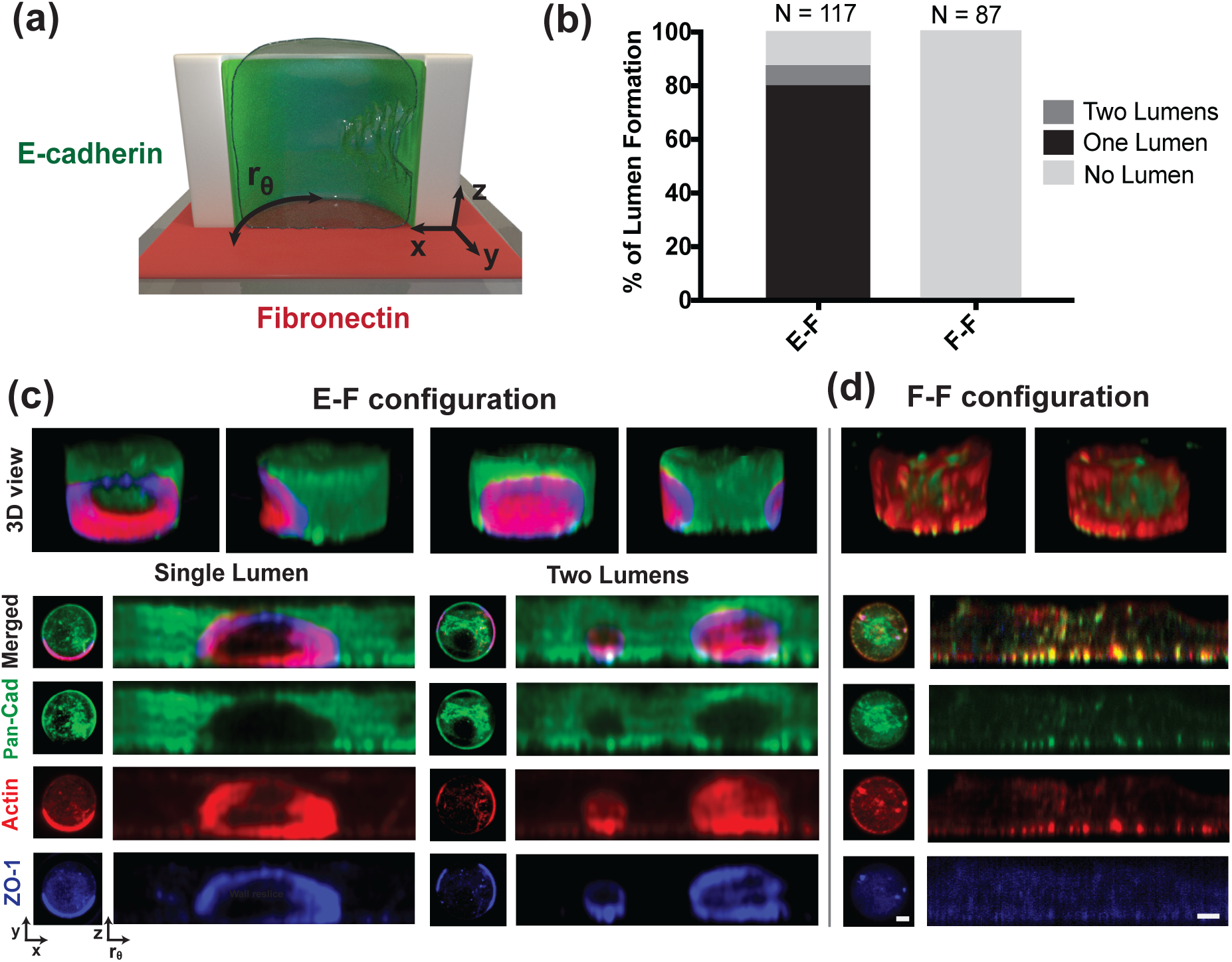
Hepatocytes apico-basal polarization in the micro-cavities. **(a)** Schematics of a single primary rat hepatocytes seeded into a micro-well coated with fibronectin at the base layer and E-cadherin on the side wall of the micro-well (E-F configuration). In F-F configuration, the microwells base and side wall are both coated with fibronectin. Here, the micro-cavities are 25 µm in diameter, 18,5 µm in height and fabricated in NOA 73. **(b)** Percentage of the hepatocytes forming lumen structure in E-F versus F-F configuration. In E-F configuration, 80% (N=117) of the hepatocytes display a lumen and 7% even show two pre-apical patches, in only 13% of the cases, cells had no specific structures. Such protein distribution was not visible for the F-F coated wells. **(c)** Example of hepatocytes with one or two lumens. Left panel: Maximum intensity projection of the different proteins along z axis. Right panel: Ortho-radial projection of the protein intensity along the micro-cavity side wall. The lumens are characterized by an ellipsoidal region depleted of cadherin (green) and by the recruitment of actin (red) and ZO-1 (blue) at the edge of this ellipsoidal region. **(d)**. Hepatocytes plated on homogeneous fibronectin coating (F/F coating) display a continuous cadherin and actin distribution with scattered ZO-1 puncta instead of the ellipsoidal protein organization observed in E/F. Scale bar **(c), (d)**: 5 µm.

### 2.4 *in situ* differential patterning with hydrogels

We now demonstrate that matrices of various stiffness can be included in the cavities in combination with protein patterning. In the absence of any modification, the Young’s moduli of the membranes are in the MPa range (11 MPa for NOA and 5.6 MPa for BIO-134) that both correspond to rigid substrate for the cells. Our approach consists in locally coating the micro-niches with softer substrates (300 Pa to 30 kPa). We successfully used glass coverslips spin-coated with a 20 µm thick layer of soft PDMS (30 kPa after curing for 24h at 80°C) to lower the stiffness of the niche base. We further tuned down the rigidity of the niches base and walls from 10 kPa to 300 Pa using hydrogels. The challenge was to ensure proper biocompatible bonding of the gels to the pit and base layers. To this end, we leveraged on the unreacted acrylate moieties and photo-initiator at the surface of the partially-cured NOA 73 pit layer (Curing conditions in **Table S2**). **Figure 6a-b** illustrates the essential steps to create a hydrogel base or niches coated with hydrogel walls (Full details in **Note S2**). The layers are immersed in a hydrogel premix deprived from any photo-initiator. Under UV exposure, the hydrogel grows from the pit surface until it fills the entire illuminated area (**Figure S2a, Movie S5-6**). The 10s illumination can be homogeneous (**Figure 6a**) or structured with PRIMO (**Figure 6b-c)**. It enable the niche functionalization with soft base layers (**Figure 6a**), soft barriers (10 to 1 µm thick) splitting the entire niche (**Figure 6b**) or soft walls of the wells (**Figure 6c**). We achieved comparable results with several hydrogels types: *i-* inert UV polymerizable hydrogels (PEGDA 700Mn, 2000Mn (Sigma), 3400Mn (ESI BIO)), *ii-* extracellular matrix hydrogels (HyStem hydrogel (EsiBio), Methacrylated Collagen I (Advance BioMatrix)), *iii-* a mix of PEGDA and thiol-containing recombinant ECMs (fibronectin and laminin) or *iv-* cellular extract (Matrigel). Hence, hydrogels of different chemical composition enables to multiplex the hydrogels coating in a single niche with collagen gel stained with FITC, around the surface of the niche, and recombinant laminin-PEGDA gel (stained with a His-Tag-DyLight 680) forming a wall in the center (**Figure 6c**).

**Figure 6.**
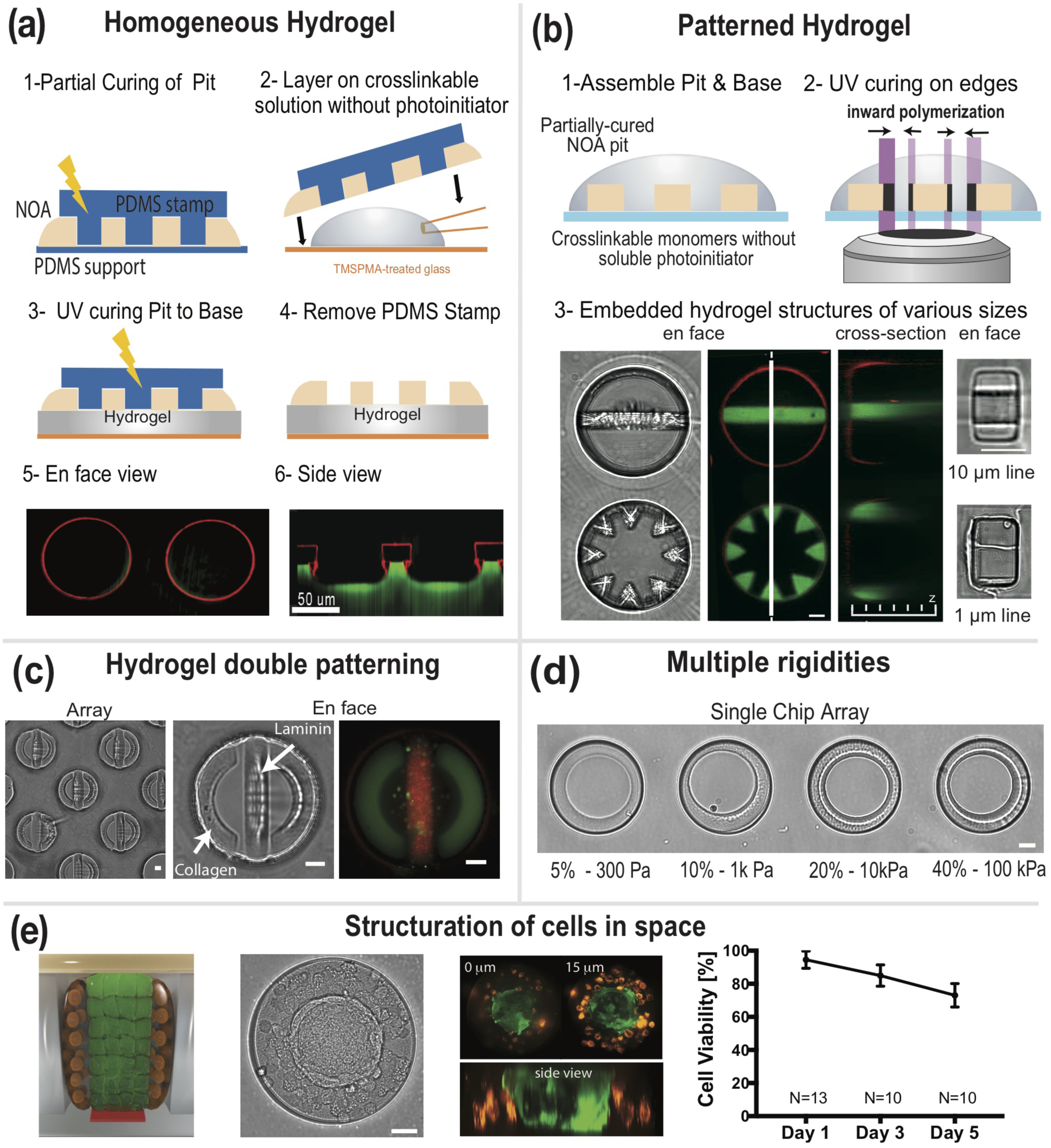
Hydrogel growth for the control of the local stiffness of the micro-environment. **(a)** Schematics of fabrication of a homogeneous hydrogel base layer covalently bonded to the pit layer **(Note S2**). A partially cured NOA 73 pit membrane is fabricated with a PDMS stamp (1). The membrane and PDMS stamp are placed together on a drop of hydrogel premix solution without photo-initiator (2) that is subsequently cured under UV (375nm) homogeneous illumination (3). Removing the PDMS stamp leaves the cured pit with a covalently bonded soft hydrogel base (4). Example of en face view (4) and side view (5) of fibronectin rhodamine (red) coated pit bounded to a 20% PEGDA hydrogel containing 0.1% FITC (green). (**b**) Schematics of *in situ* growth of patterned hydrogel inside of the micro-niche. The hydrogel premix placed on partially cured microwells (1) is locally cross-linked from the micro-well edges by structured UV illumination obtained with PRIMO (2). Representative examples of locally cross-linked hydrogel structures across the microwells of various features and size (3). The bright field and fluorescence images show an array of two microwells of 60 µm diameter with two different hydrogel patterns (20% PEGDA with 0.1% FITC). The gel is cross-linked all along the pit height (20 µm) from the pit edge inward (3 – cross-section). The bright field images from rectangular microwells (20 µm x 10 µm x 10 µm) display hydrogel stripes from 10 µm to 1 µm thick that are stabilized by the NOA 73 pit backbone (3-left). **(c)** Bright field and fluorescent images of a double ECM hydrogels pattern composed of a stripe of recombinant laminin-base gel (red from His-Tag conjugated DyLight 680) and a ring of Methacrylated collagen gels (green from FITC) obtained by successive crosslinking of the two ECM gels (**d**) Bright field images of a single chip composed of four microwells with hydrogel rings of different rigidity (0.3, 1, 10, 100 kPa) obtained by successive incubation with increasing concentration of PEGDA (5, 10, 20, 40 wt%). The storage young modulus for each PEGDA concentration has been measured for bulk hydrogel. **(e)** Structuration of different cell types in space. The phase contrast and fluorescence images show the live imaging of primary tumor HN cells (steadily expressing GFP, green) pre-formed as spheroids and seeded into the centre of microwells (200 µm diameter, 60 µm height) coated with fibronectin (red on the design illustration) surrounded by metastatic HN cells (steadily expressing RFP, orange) encapsulated inside a methacrylated collagen-PEGDA hydrogel. We tested the viability of the co-culture over a week. Scale bars (**b**), (**c**), (**d**), (**e**): 10 µm.

Likewise, **Figure 6d** shows circular wells coated with PEGDA of various stiffness (300 Pa to 10 kPa,) created by sequential polymerization from PEGDA solution at increasing concentrations (**Figure S3b**) on the same chip. The 10s exposure time per field of view (**Table S1**) enables thousands of niches to be constructed in an hour (**Movie S7)**.

The successful testing of this wide range of compatible matrix proteins demonstrates that stiffness can be adjusted independently, without sacrificing the biochemical cues. Taken together, we demonstrate that the ability to control the local rigidity does not compromise the ability to structure the niche in 3D with different ECM proteins. In our approach, the base and pit (NOA 73) act as rigid backbones that ensure the structural integrity of the niche, while the hydrogel coating locally modulates the stiffness felt by the cell. Since swelling is proportional to the total volume of hydrogels, the minimal volume of hydrogel we use to create the features largely minimizes the absolute changes of the dimensions of the hydrogel features. Moreover, the hydrogel layers can be preserved for at least 6 months by freeze-drying (**Figure S3c**).

Cells can also be embedded in the polymerized hydrogel to structure cell co-culture in space. We grew a layer (50 microns) of PEGDA (3400 Da)-methacrylated collagen I hydrogel (**Figure 6e**) on the walls of the wells (200 µm diameter and 60 µm height) with embedded metastatic Head and Neck (HN) tumor cells. We tested their viability over a week (Figure S2). We then seeded epithelial HN tumor cells, stably expressing GFP in the center of the hydrogel doughnut (**Figure 6e**). We successfully grew the co-culture that displays rearrangements of the cells from the aggregate and the gel (N=5) (**Movie S8, Figure S4**).

### 2.5 Additional control over topographical cues

Lastly, the local topography of the niche can be controlled in combination with biochemical and rheological cues. Micro-textured surfaces are now commonly used in various fields of cell biology, either as mechanical cues or as mechanical sensors (e.g., Traction Force Microscopy).^[34,35]^ Such surfaces can easily be combined with our toolbox. We used homogeneously textured bases with nano/micro topographical features (groves, pillars, lenses) created by UV curing or hot embossing (**Figure 7a-b**, Nanosurface Biomedical (USA) or home-designed). Grooves are the only topographical features that we incorporated along the vertical axis of the pit due to fabrication constraints (**Figure S5**). Topographical features in the niche does not hamper the direct patterning of protein on either the wall or the base. **Figure 7c** shows individual niches (60µm diameter, 30µm height) coated with laminin on their sides were deposited on a base layer textured with BIO-134 micro-pillars (2 µm diameter, 2 µm in height) and coated with fibronectin. Clusters of patient derived HN cells were also grown in such micro-niches. Topographical features also allow hydrogel growth as demonstrated in **Figure 7d**.

**Figure 7.**
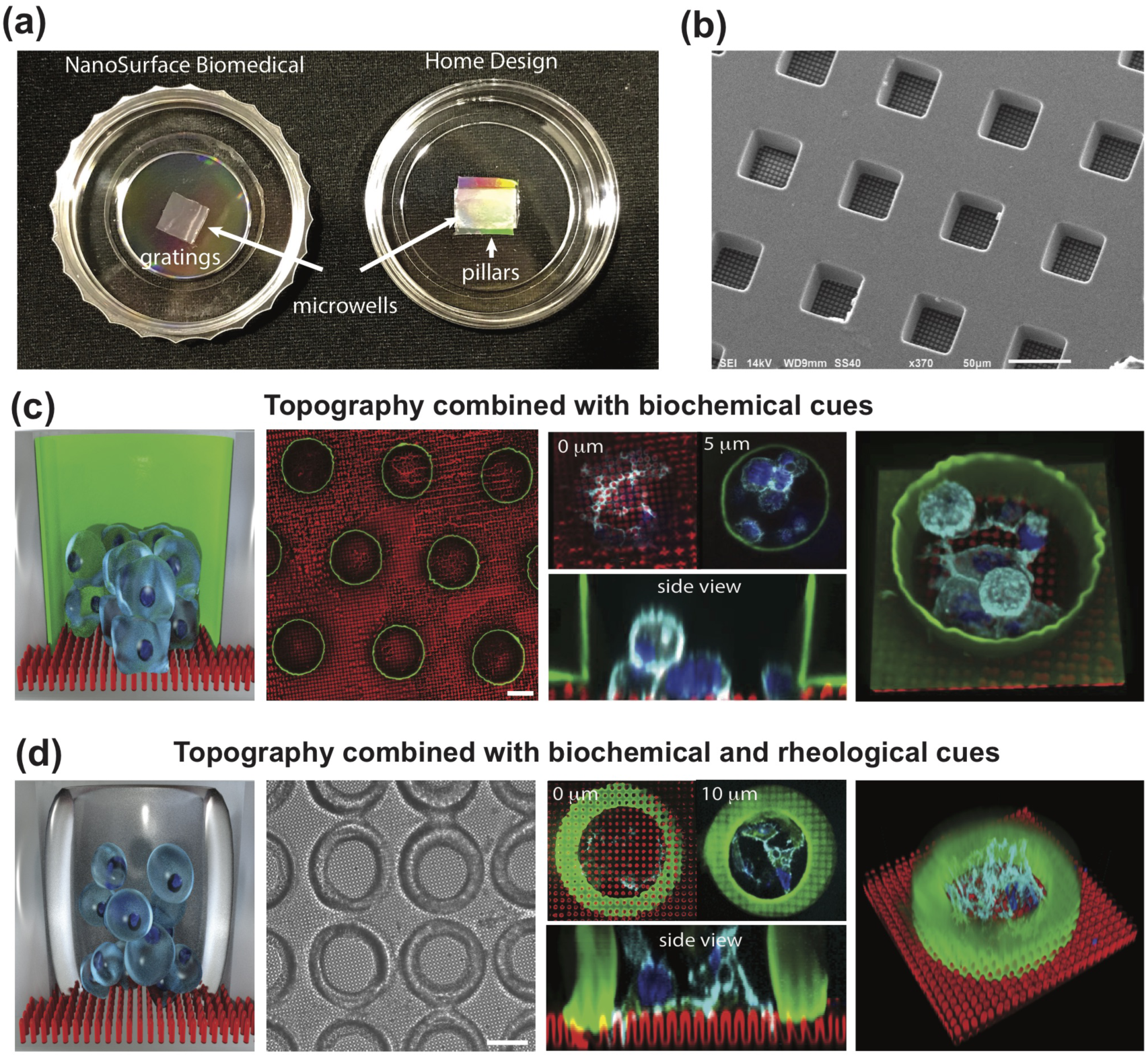
Control of the topography at the base of the micro-niche combined with increasing complexity of environmental cues in 3D. **(a)** Micro-textured surfaces combined with NOA 73 pit layer for a commercially available Nanopatterned dish from Nanosurface Biomedical (left) and for a “home-made” dish with pillars (right) **(b)** SEM images of NOA 73 microwells (30µm square x 40µm height) assembled with nano-gratings (800nm width x 600nm height with 800nm gap) from the NanoSurface Biomedical dish (Bottom left) and assembled with a base layer of “home designed” pillars (2 µm diameter and height) for the home design dish (bottom right). **(c)** Combining topographical cues and biochemical cues at cell aggregate level by addition of a micro-pillars base layer. Patient-derived head and neck cancer cell aggregates (HN cells) were grown into 60 µm diameter and 30 µm height micro-niches with Laminin (green) homogeneously coated on the pit walls, and fibronectin (red) coated on the NOA 73 micro-pillars (2 μm height and diameter) were visualized by staining actin (cyan) and nuclei (blue). **(d)** Additional rheological cues were combined to the previous niche. Recombinant Laminin-base gel with 0.1% FITC (green) was cross-linked as a ring on the side of 60 µm diameter microwells placed on MyPolymer 134 pillars (5 μm height, 2 μm diameter) coated with fibronectin (red). Patient-derived HN cell aggregates were culture for one day into this microenvironment exposing the cells to a softer rigidity E∼2kPa on the side of the micro-niche. HN cell shapes were visualized by staining actin (cyan) and nuclei (blue). Scale bar **(c), (d)**: 30 µm

## 3 Discussion

Microwells have long been recognized as a good way to isolate individual cells or aggregates ^[36,37]^. A large number of technological approaches (eg: etching, embossing, fusing and printing) can be used to create micro-cavities. They all rely on creating pit arrays in a supporting substrate. Subsequent uniform coating with a single protein type (usually ECM) fosters cell adhesion and turns the pits into micro Petri dishes. Our methods using a stacked deposition of UV reactive membranes coupled to UV protein printing enables to turn the micro-dishes into micro-niches with differential and adjustable modulation in 3D of rheological, topographical and biochemical cues presented to the cells. Different combinations are obtained depending on the sequence of membrane lamination and bio-functionalization. **Table S1** summarizes the time order of the various protocols that have to be applied to obtain a given combination of cues. The approach uses exclusively commercially available materials that can readily be adopted by other labs. Increasing the complexity of the niches does not involve extra steps nor additional complexity in the niche assembly but relies exclusively on the nature of the protein/hydrogel used and the preformed structures of the various layers.

The bio-functionalization of the niches can be done in bulk for differential homogeneous coating on the pit and bottom layer. It requires the use of a UV protein printing system (here PRIMO) for more intricate patterns. The increase in preparation time is minimal due to the ability of multi-well printing by bright field illumination. Functionalizing 1000 identical wells requires around 1 hour (**Figure 3, Movie S4**).

All in all, the method combines standardized cell culture conditions with well-defined biochemical and biophysical parameters in 3D. Thousand of niches can be fabricated within an hour and are compatible with high-resolution observation of cell behavior. We believe that taken together these characteristics make our approach a tool of choice to study environmental sensitivity at the single cell/single aggregate level.

### Experimental section

#### Fabrication of PDMS mold and pit layers

Negative PDMS (polydimethylsiloxane) molds with different sizes and shapes of pillar features were fabricated by standard lithography techniques (**Note S1**). Briefly, silicon wafers with desired patterns were fabricated in-house in SU8-3050 using classical lithography technics. Then, the wafer was silanized (Trichloro-(1H,1H,2H,2H-perfluorooctyl)silane, Sigma-Aldrich, MO) in a vacuum jar for 6 h before pouring a 1:10 mix of PDMS elastomer and curing agents (Sylgard 184, Dow corning, MI). The PDMS liquid mixture was cured at 80°C overnight. Pit layers were made by capillarity infiltration of UV curable polymers, NOA 73 (Norland Products Cranbury, NJ) or BIO-134 (MYPolymer, Israel) into the PDMS mold placed against glass coverslips, glass-bottom dishes (Iwaki, AGC Techno Glass, Japan), plastic-bottom dishes (Ibidi, Germany), or flow chambers (Ibidi, Germany). Subsequently, the polymers were cured by UV illumination (365 nm, UV-KUB 9, KLOE, France) under conditions listed in **Table S2**. Final micro-well structures were checked by the scanning electron microscope (JSM-6010LV with Au/Pd sputter coater JFC-1600, JEOL, Japan).

#### Lamination and functionalization of membranes with various properties

Pit layers and based layers were bio-functionalized, separately or together, using three ways of membrane assembly and four functionalization protocols (**Table S1** and **Note S2)**. Pits were removed and transferred by tweezers from the template surface onto base surfaces, which can be glass/plastic bottom, polymeric pillars or deformable substrates. Membranes were pressed gently to ensure full contact. For base layers made of pillars or hydrogels, partially cured pit layers were required to provide good adhesiveness. Stacking of two pit layers (**Figure 2c**) was achieved using microscope stage equipped with a positioner (Siskiyou, OR).

#### Multi-protein printing by Light-Induced Molecular Adsorption

After 3 min of oxygen plasma treatment, NOA 73 polymer pits, were coated with PLL-g-PEG (PLL(20)-g[3.5]-PEG(5), SuSoS AG, Dübendorf, Switzerland) at 100 µg.mL^-1^ in PBS for 1h. PRIMO system (Alveole, France) mounted on an inverted microscope (Nikon Eclipse Ti-E, Japan) equipped with a motorized scanning stage (Physik Instrumente, Germany) was used to form the DMD-based UV pattern at the focal plane of the microscope. After alignment of pit with the UV pattern by using the Leonardo software (Alveole, France) and moving the scanning stage, a solution of photoinitiator (PLPP, Alveole, France) was incubated on the sample. The UV-activated photoinitiator molecules locally cleaved the PEG chains in a UV dose dependent manner. After rinsing the PLPP solution, the microwells were incubated with a fibronectin protein solution (Rhodamine Fibronectin, Cytoskeleton, Inc., 50µg.mL^-1^) which adsorbs to the UV-exposed areas. For two proteins patterning, after rinsing the first fibronectin protein solution with PBS, Fibrinogen Alexa 488 (ThermoFisher Scientific, 50µg.mL^-1^) was patterned using the same procedure. The two patterns of protein are aligned with 500 nm precision corresponding to the scanning stage precision. (See also **Note S2**)

#### Crosslinking of hydrogel patterns without photo-initiator

Partially cured NOA 73 pit layer were prepared with capillarity filling of a PDMS stamp (Round shape pillars, 60µm) by curing under UV 365nm light at 20mW/cm^2^ power (10%) for 30 s. Subsequently, hydrogel precursors were directly placed on top of the surface to fill the NOA pit layer without any washing steps. 20wt% Polyethylene glycol diacrylate (PEGDA, Mw 700, Sigma-Aldrich, MO) and 5wt% thiol-containing fibronectin/laminin or methacrylate-modified collagen (Advanced BioMatrix, CA) were mixed with PBS pH 7.4 to establish hydrogel precursor solutions. Upon precursor filled wells, PRIMO system was used with 375nm UV laser (4mW/cm2) and defined digital micromirror device (DMD) patterns to locally crosslink the PEGDA hydrogel. The region of the projected pattern must overlap the NOA/precursor interface to activate the unreacted photo-reactive groups for initiation of free-radical mediated PEGDA gelation. Reaction were conducted at ambient conditions. The time of laser irradiation was 2 sec. Patterned gelation were visible from the live bright field imaging. (See also **Note S2**) After hydrogels being patterned, the micro-dish was subject to UV 365nm at 5% power 1 min (cytocompatible energy) to fully cure the substrates and avoid side effects from half-cured NOA. Fluorescent hydrogels were made with 0.1% FITC dye. Freeze dried microwells with hydrogel micro-patterns (shown in **Figure S3**) were immersed in PBS overnight and then frozen at −80 followed by lyopholization using freeze dryer.

#### Rheological measurement

The stiffness of the hydrogel were measured by a Physica MCR 501 rheometer using Rheoplus software (Antor Paar, Graz, Austria). PEGDA hydrogels were fully cured with 0.1wt% photoinitiator (lithium phenyl-2,4,6-trimethylbenzoylphosphinate) and swollen in PBS overnight. The swollen hydrogels were placed between 8 mm diameter parallel plates and rheometer stage. Dynamic frequency sweeps were performed at a constant stain at 1% with the angular frequency was varied from 0.2 to 10 rad/s. All samples were measured at 37°C to simulate physiological conditions. Storage Young modulus was derived from storage shear modulus by assuming Poisson’s ratio at 0.5.

#### Cell culture

Human foreskin fibroblasts (HFF) and HaCat were culture in DMEM (High-glucose, ThermoFisher) supplemented with 10% fetal bovine serum (FBS, Gibco, CA) and 1% penicillin/streptomycin and used at passage 5-10. All cells were trypsinized with 1x TrypLE Express (Gibco, CA) before seeding to the micro-dishes. Cells were seeded with a 20µl droplet (1 million cells per ml) to cover the 5 mm^2^ membrane surface. Single cells of HFF and Hacat were spin on the microwells by centrifugation at 500 rpm for 2 min. After 30 min, dishes were gently rinsed to remove excessive cells outside the pit. The petri dish was subsequently replenished with fresh culture medium for another 8 hours before fixation. Patient-derived head and neck (HN) tumor cells, primary and metastatic cells (wild type), were obtained from Genome Institute of Singapore (GIS). GFP-expressing primary tumor cells and RFP-expressing metastatic HN cells were also received from GIS. Those cells were maintained in RPMI media (Gibco, CA) supplemented with FBS, 1x Glutamax (Gibco, CA), and 1% penicillin/streptomycin. They were used between passage numbers 4 to 30. HN cell aggregates could fall into 60-300 µm pits by gravity. The surface of the pit was incubated with 0.2% (w/v) Pluronic acid (Pluronic F-127, Sigma-Aldrich, MO) at the final step. Hepatocytes were isolated from male Wistar rats by a two-step in situ collagenase perfusion method. Freshly isolated rat hepatocytes (0.5 million) were seeded onto the microwells and cultured in 2 ml of William’s E culture medium supplemented with 2 mM L-Glutamine, 1 mg/ml BSA, 0.3 µg/ml of insulin, 100 nM dexamethasone, 50 µg/ml linoleic acid, 100 units/ml penicillin, and 100 mg/ml streptomycin (Sigma-Aldrich, MO). After a 30-minute incubation, the system was rinsed with William E medium to remove cells that did not trap in the micro-niche. The petri dish was subsequently replenished with fresh culture medium for another 24 hours before fixation.

#### ECM hydrogels patterning with encapsulated HN cells and cell viability

Precursor solution was made by dissolving 5wt% PEGDA (Mn 3400, ESI BIO, CA) in CO2-independent media (Gibco, CA) and 4mg/ml methacrylate-modified collagen (Advanced BioMatrix, CA) at 1:1 ratio gently mixed on ice. Primary wild-type and stably-expressing RFP HN cells were harvested from culture flask and centrifugation at 500 rpm for 3 min to form dense cell pallets. Cells were suspended in 10 µl precursor solutions and added onto the half-cured NOA wells, followed by crosslinking with PRIMO to form hydrogel patterns. Encapsulated cells viability assay was carried out over 5 days and compared to 2D culture conditions. Wild-type HN cells were trypsinized and seeded onto a tissue culture dish (2D) or encapsulated in Collagen-PEGDA precursor solution at 2 million cells per ml, followed by staining with LIVE/ DEAD Cell Imaging Kit (Life Technologies) after 1,3 and 5 days. Conventional (2D) or confocal (3D) fluorescence imaging was used to assess the proportion of live (green) versus dead (red) cells by manual counting on three independent samples (n=3 minimum for each conditions: day1, day3, day5 (3D) and 2D).

#### Spheroid formation

Spheroids of primary HN cells were formed by seeding and spindown 1 × 10^6^ cells onto a 6-well plate of 400µm AggreWell (StemCells Technologies), previously casted in agar to avoid cells attachment. This method allows the formation of 4700 spheroids of uniform 100µm size per 6-well plate. Spheroids were collected after overnight incubation and seeded onto the functionalized pit layer.

#### Immunostaining

Cells were fixed with 4% paraformaldehyde (ThermoFisher) for 15min. Cells were permeabilized with 0.5% Triton-X100 (v/v) for 10 min. Subsequently, the samples were blocked with blocking buffer (PBS containing 2% BSA and 0.1 M glycine) for 30 min. For primary antibody labeling, rabbit polyclonal anti-ZO-1 (Life Technology, 61-7300,1:100) and mouse monoclonal anti-Pan-Candherin (Sigma, C1821, 1:200) were incubated overnight at 4°C. For secondary antibodies, we used Alexa Fluor 546 Donkey Anti-Rabbit IgG (Thermo Fisher, A10040, 1:200) and Alexa Fluor 488 Donkey Anti-Mouse IgG (Thermo Fisher, R37114, 1:200). Cells were stained with phalloidin-Atto 633 (Atto-tec, Germany, 1:200) to visualize F-actin and DAPI to visualize the nucleus (Cell Signaling, MA) for 1h at RT.

#### Microscopy and image analysis

Micro-niches were imaged using a confocal scanner unit (CSU W1, Yogogawa, Japan) mounted on an inverted microscope (Nikon Eclipse Ti-E) equipped with a motorized scanning stage (Physik Instrumente, Germany), a digital CMOS camera (ORCA-Flash4.0, Hamamatsu, Japan) and a 60× oil-immersion objective. Metamorph software (Molecular Devices) was used to control the *x-y-z* stage positions.

#### Image Analysis and Data processing

Matlab R2015b (Mathworks) was used to analyze and process the image stacks of the cells on different 3D geometrical patterns. The acquired images were aligned by using the fluorescent micro-well contour in DAPI channel and stacked together for each Z plane. The average spatial distribution of actin in 3D was obtained by calculating the average intensity of each pixel over the stack (**Figure 4**). DAPI channel stacks were filtered using thresholding and segmentation to automatically detect the position of nucleus for each cell in 3D. Data were further projected along the xy or xz axes to obtain the distributions represented in **Figure 4**. After data processing with Matlab, 3D stacks and 3D average actin intensity were reconstructed, visualized and recorded using Icy.^[38]^ For **Figure 5**, Z and ortho-radial projections of the hepatocytes image stacks in E-F and F-F conditions were obtained using Image J.

#### Statistical analysis

All results were expressed as mean ± standard error with n=3 mininmum for all experiments and 3 independant samples. Statistical analysis was performed with Prism software and one-way analysis of variance (ANOVA). Significant differences were indicated by (P < 0.05).

