## Supplementary Materials for "A new approach to design artificial 3D micro-niches with combined chemical, topographical and rheological cues"


| | |
|---|---|
| **Supplementary Figure 1** | Steps during cells seeding on the microwells |
| **Supplementary Figure 2** | Viability of the cells encapsulated in a patterned hydrogel inside the micro-niches cross-linked with PRIMO system |
| **Supplementary Figure 3** | Hydrogel growth kinetics, rigidities measurement for different PEGDA concentration and preservation of the cross-linked gel on the pit by freeze drying. |
| **Supplementary Figure 4** | Examples of spatially structured co-culture with HN GFP-labelled cell spheroid surrounded by a HN RFP-labelled cells encapsulated in a hydrogel ring. |
| **Supplementary Figure 5** | Vertically textured pit with grooves |
| **Supplementary Movies** | Legends for the supplementary movies 1 – 8 |
| **Supplementary Table 1** | Series of functionalization protocols to create any combination of environmental cues following the fabrication. |
| **Supplementary Table 2** | Curing conditions for the polymers and hydrogels |
| **Supplementary Table 3** | Characteristics of the illumination settings and size of the patterned niches |
| **Supplementary Note 1** | Fabrication of pit layer membrane by soft lithography |
| **Supplementary Note 2** | Protocols for Supplementary Table 1: Assembly and functionalization of the micro-niches |

### Supplementary Figures:

#### Supplementary Figure 1:

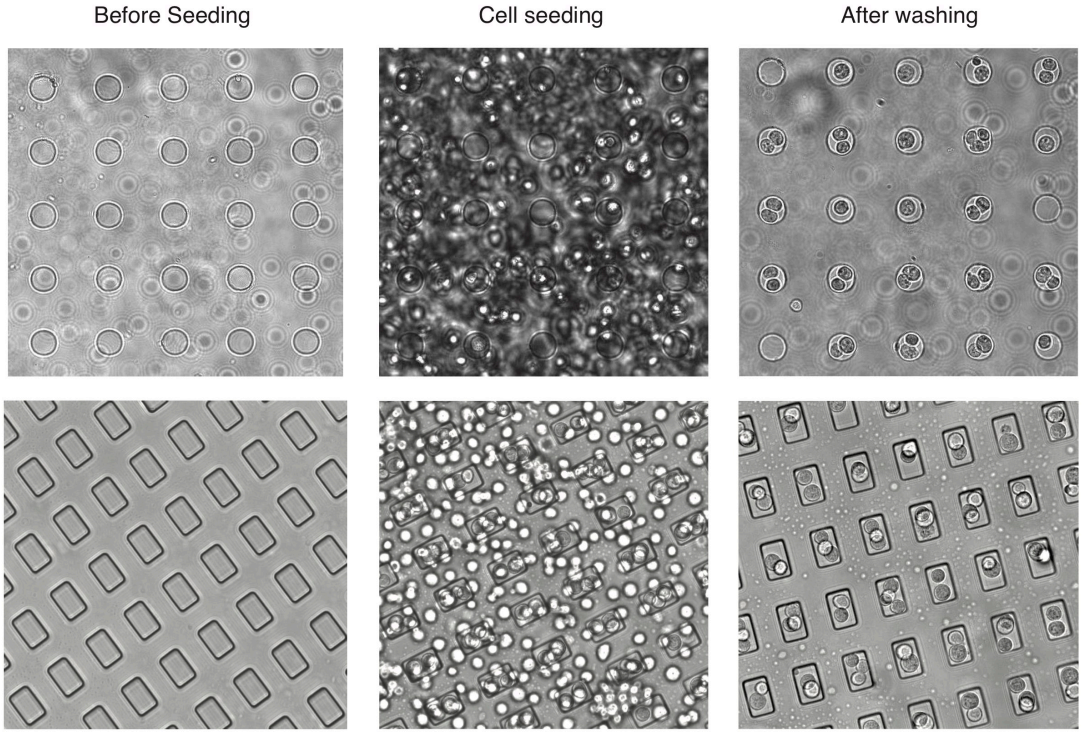


**Supplementary Figure 1:** Steps during cell seeding in the microwells of different shapes. The PBS is first replaced by warm media. The media is then removed without drying and the cells are seeded with a density from 100-200 cells/µL on the pit layer. The cells fall by gravity into the microwells and are incubate for 30min -1h depending on the cell type. Alternatively, the cells can be spin down onto the microwells. The excess of cells is washed away with warm media. If the cells seeding density is not sufficient, the seeding procedure can be can be repeated by following the previous steps.

**Supplementary Figure 2:**

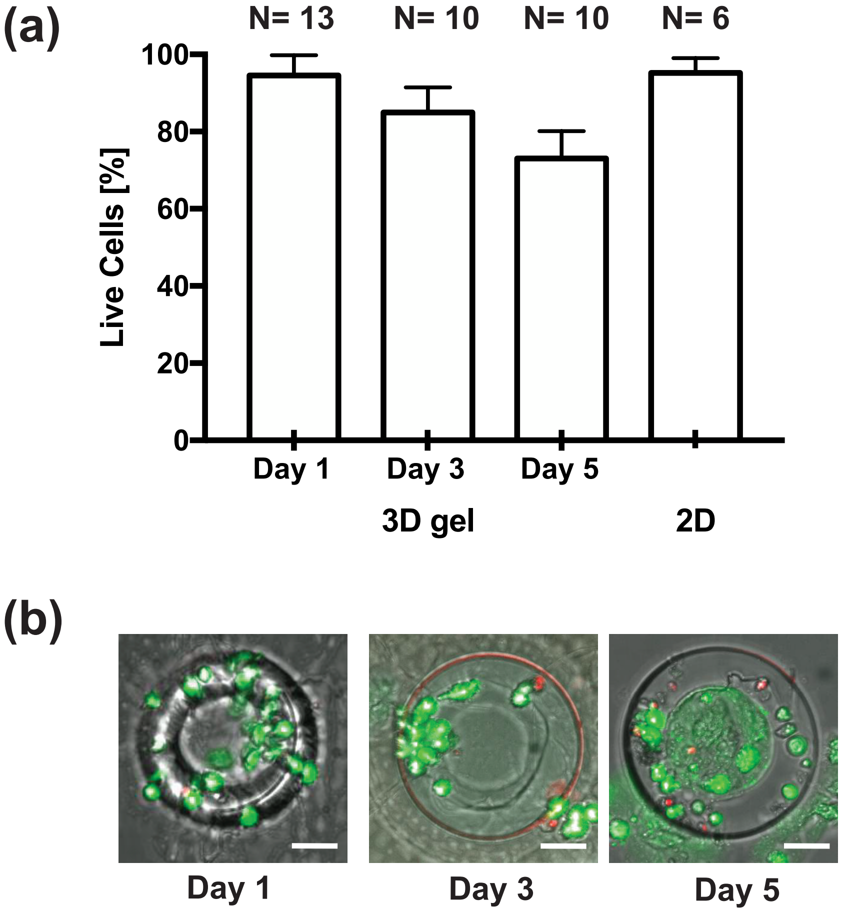


**Supplementary Figure 2: (a)** Viability assay of wild-type primary HN cells encapsulated in a ring of collagen-PEGDA (5% PEGDA (Mn=3400)), 2mg/ml metacrylated collagen) hydrogel inside the micro-niches cross-linked with PRIMO system. After one day of culture no significant difference in viability can be seen between 2D and 3D culture. After 5 days of culture the viability is around 70%. The cells migrating out of the hydrogel layer after 3 days can account for the observed drop. **(b)** Representative images of live/dead staining superimpose with phase contrast images of the wild type primary HN cells after 1,3 and 5 day of encapsulation in a methacrylated collagen-PEGDA hydrogel ring. Scale bar **(b)**: 20 µm

**Supplementary Figure 3:**

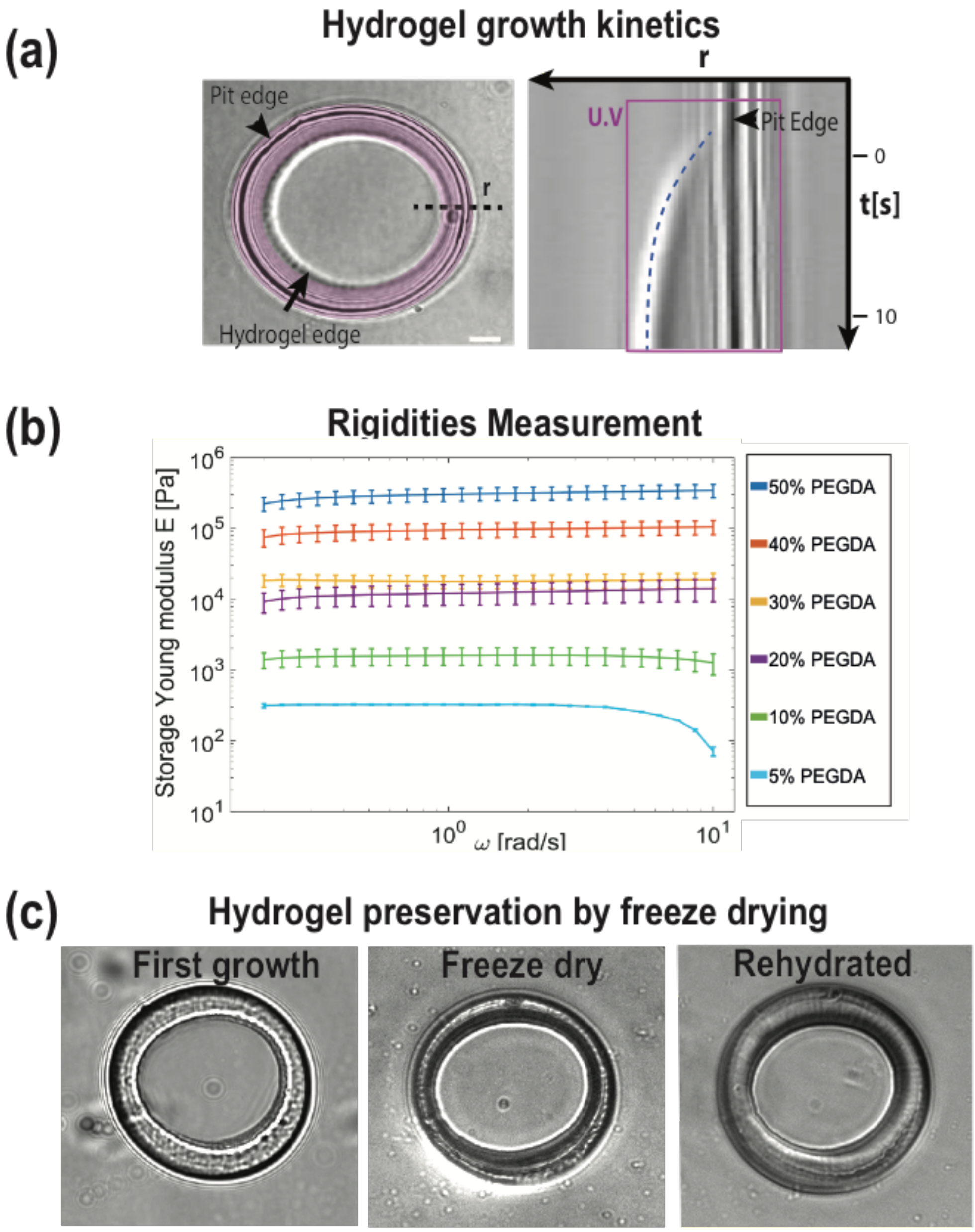


**Supplementary Figure 3: (a)** Kymograph displaying the growth kinetic of a PEGDA (20%) hydrogel ring along the dashed line across the 60 µm diameter micro-well. The growth starts from the pit edge to the edge of the UV illuminated region (in purple). This effect is due to the use of the photo-initiator contained in the partially cured NOA pit. **(b)** The different rigidities (0.3, 1, 10, 100 kPa) are measured for each PEGDA concentration (5, 10, 20, 30 40 and 50 wt%, Mn=700) for bulk hydrogel with a rheometer. **(c)** Phase contrast images of microwells (NOA 73, 60 µm diameter and 30 µm height) with 20% PEGDA hydrogel ring patterns after immersion in PBS overnight, during freeze-drying and after a second rehydration with PBS.

**Supplementary Figure 4:**

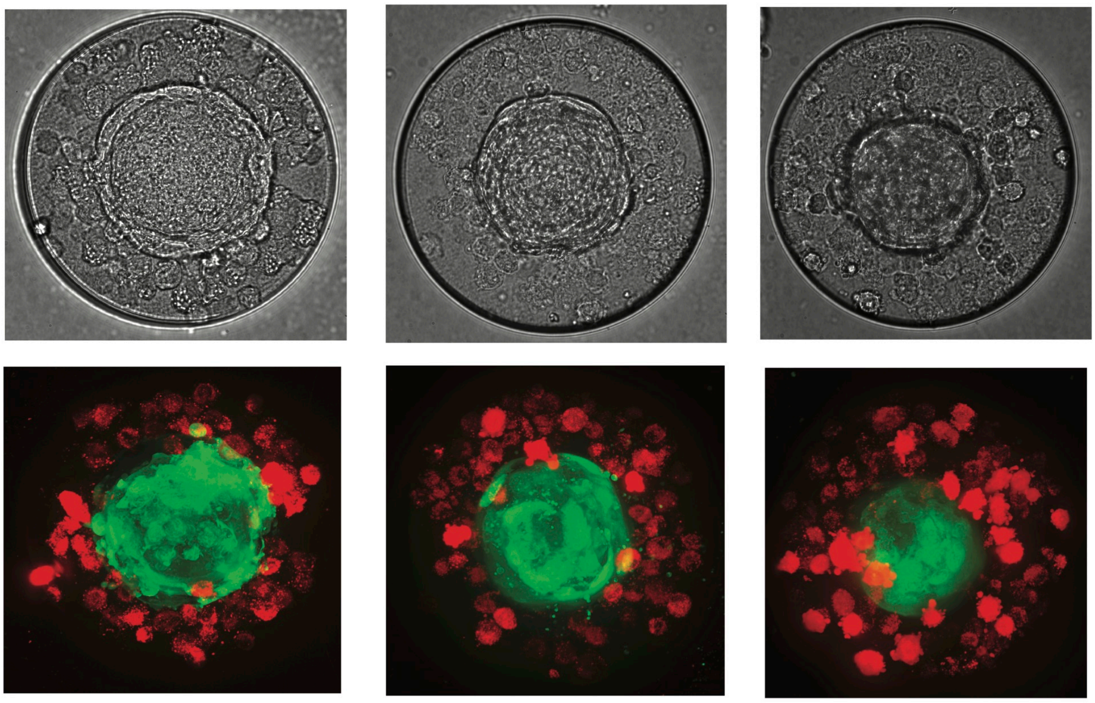

**Supplementary Figure 4:** Examples of different spatially structured co-culture with HN cells stably expressing GFP grown as spheroids surrounded by a ring of HN cells stably expressing RFP encapsulated in a methacrylated collagen-PEGDA hydrogel (5% PEGDA (Mn=3400)), 2mg/ml metacrylated collagen I) ring bonded to the wall of the niche (200 µm diameter and 60 µm deep microwells, NOA73). **Top:** phase contrast images of the co-culture. **Bottom:** Maximum intensity projection of the live confocal imaging.

**Supplementary Figure 5:**

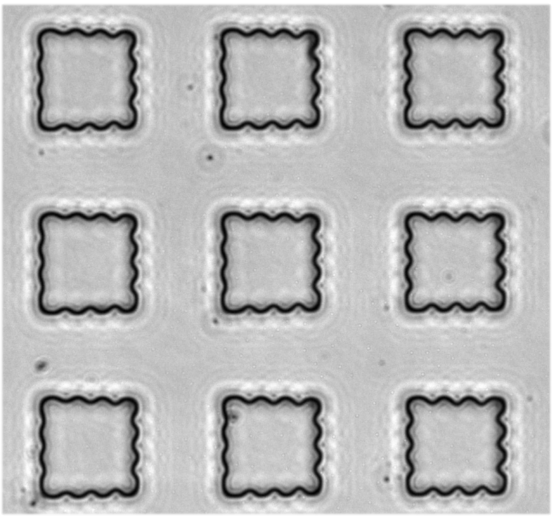

**Supplementary Figure 5:** Vertically textured pit with grooves features. Phase contrast images of microwells (30 µm cubes) with grooves of 3 µm separated by 5 µm gap.

#### Supplementary Movies

**Supplementary Movie 1:** Rotating 3D visualization of a hemispheric pit (50µm diameter and 30 µm height) of NOA 73 patterned with a fibronectin rhodamine checkboard (red).

**Supplementary Movie 2:** Rotating visualization of the average actin intensity in 3D (crossbow N=22, triangle N=28, cross N=34, plain circles N=25) showing the hepatocytes cytoskeleton 3D structure on different geometrical shapes.

**Supplementary Movie 3:** Combined confocal images of 9 hepatocytes cells on the different 3D geometrical patterns (crowssbow, triangle, cross, circles) and 9 HaCat cells on crossbow pattern.

**Supplementary Movie 4:** Rotating 3D visualization of one hepatocytes in the well in F-F configuration and E-F configuration with single and double lumen. The hepatocytes are stained for Pan-cadherin (green), Actin (red) and ZO-1 (blue).

**Supplementary Movie 5:** Time-lapse imaging of *in situ* growth of PEGDA hydrogels inside a 60 µm diameter pit. The growth starts from the pit edge to the edge of the UV illuminated region (ring shape). This effect is due to the use of the photo-initiator contained in the partially cured NOA pit to locally grow a hydrogel layer from the micro-niche surface. The growth completes in 10 sec.

**Supplementary Movie 6:** Demonstration of *in situ* growth of PEGDA hydrogels from the pit. Microwells at the left hand side were filled with PBS only, while at the right hand side they were filled with PEGDA solution (Mw700, 20w% in PBS, without any photoinitiators). Both wells were subject to the same stripe of UV illumination. PEGDA hydrogels can be spatially crosslinked without photoinitiator inside a microwell.

**Supplementary Movie 7:** Moving field of view demonstrates the scalability of the technique to create arrays of microwells patterned with hydrogels.

**Supplementary Movie 8:** Time-lapse imaging of a microniche with co-culture for 24 hr. Primary HN cell spheroid adhered on the bottom coated with fibronectin, and neighboring metastatic HN cells encapsulated in a ring-shape collagen gels.

### Supplementary Table 1:

<table>
<tr><td colspan="4" rowspan="4"></td><td colspan="5">BASE (B) Layer</td></tr>
<tr><td colspan="3">Homogeneous biofunctionalization</td><td colspan="2">Patterned biofunctionalization</td></tr>
<tr><td colspan="2">GPa - 15kPa</td><td>Hydrogel (10kPa-100Pa)</td><td colspan="2">GPa - 15kPa</td></tr>
<tr><td>I</td><td>II</td><td>I</td><td>I</td><td>II</td></tr>
<tr><td rowspan="4">PIT (P) Layer</td><td rowspan="2">Homogeneous biofunctionalization</td><td>10MPa</td><td>I<br>II<br>III<br>IV</td><td>1. Protocol Base : A<br>2. Protocol Pit : A<br>3. Assembly 3</td><td>1. Protocol Base : A<br>2. Protocol Pit : A<br>3. Assembly 3</td><td>1. Protocol Base and Pit : D<br>2. Protocol Pit : A</td><td>1. Assembly 1<br>2. Protocol Base and Pit : B</td><td>1. Assembly 2<br>2. Protocol Base and Pit : B</td></tr>
<tr><td>Hydrogel (100 Pa- 10kPa )</td><td>I<br>II<br>III<br>IV</td><td>1. Protocol Base : A<br>2. Assembly 1<br>3. Protocol Pit : C</td><td>1. Protocol Base : A<br>2. Assembly 2<br>3. Protocol Pit : C</td><td>1.Protocol Base and Pit : D (Special curing condition required)<br>2. Protocol Pit : C</td><td>1. Assembly 1<br>2. Protocol Base : B<br>3. Protocol Pit : C</td><td>1. Assembly 2<br>2. Protocol Base : B<br>3. Protocol Pit : C</td></tr>
<tr><td rowspan="2">Patterned biofunctionalization</td><td>10MPa</td><td>I<br>II<br>III<br>IV</td><td>1. Assembly 1<br>2. Protocol Base and Pit : B</td><td>1. Assembly 2<br>2. Protocol Base and Pit : B</td><td>1. Protocol Base and Pit: D<br>2. Protocol Pit : B</td><td>1. Assembly 1<br>2. Protocol Base and Pit : B</td><td>1. Assembly 2<br>2. Protocol Base and Pit : B</td></tr>
<tr><td>Hydrogel (100 Pa- 10kPa )</td><td>I<br>II<br>III<br>IV</td><td>1. Protocol Base : A<br>2. Assembly 1<br>3. Protocol Pit : C</td><td>1. Protocol Base : A<br>2. Assembly 2<br>3. Protocol Pit : C</td><td>1. Protocol Base and Pit : D (Special curing condition required)<br>2. Protocol Pit : C</td><td>1. Assembly 1<br>2. Protocol Base : B<br>3. Protocol Pit : C</td><td>1. Assembly 2<br>2. Protocol Base : B<br>3. Protocol Pit : C</td></tr>
</table>

**Supplementary Table 1:** Series of functionalization protocols to create any combination of environmental cues with the pit and base layer. Biochemical cues are in red (homogeneous or patterned coating), rheological cues are in maroon (different ranges of stiffness corresponding to different materials GPa to 10 kPa for Glass or plastic and 10kPa-100 Pa for ECM gel or PEG hydrogels), topographical cues are in Blue (I- Flat, II-Microstructures, III-Shape, IV-Confinement). The order of assembly and functionalization of the pit and the base are specified for each combination. The detailed functionalization protocols A, B, C and D as well as the assembly types 1, 2 and 3 are described in **Supplementary Note 2**. Special curing conditions can be found in **Supplementary Table 2**.

#### Supplementary Table 2:

<table>
<tr><td>Partially cured NOA 73 pit polymer</td><td colspan="2">0.5 $J/cm^2$ to 1.2 $J/cm^2$ 365nm</td></tr>
<tr><td>Fully cured NOA 73 pit polymer</td><td colspan="2">4$J/cm^2$ at 365nm</td></tr>
<tr><td>Fully cured BIO-134 pit polymer<br>*Immersed in water or argon gas</td><td colspan="2">1440$J/cm^2$ at 365nm</td></tr>
<tr><td>Hydrogel curing time with homogeneous illumination</td><td colspan="2">0.5 $J/cm^2$ at 365nm</td></tr>
<tr><td rowspan="4">Hydrogel curing time with PRIMO</td><td>5% PEGDA<br>HyStem hydrogel<br>Type I methacrylated collagen</td><td>0.9 $J/cm^2$ at 375nm</td></tr>
<tr><td>10% PEGDA</td><td>0.5 $J/cm^2$ at 375nm</td></tr>
<tr><td>20% PEGDA</td><td>0.3 $J/cm^2$ at 375nm</td></tr>
<tr><td>40% PEGDA</td><td>0.15 $J/cm^2$ at 375nm</td></tr>
<tr><td>Special condition for the pit and hydrogel curing in Protocol D<br>(if followed by Protocol C for the pit)</td><td colspan="2">NOA 73 Pit layer cured with 0.5 $J/cm^2$ at 365nm (step 4 of Protocol D)<br>Hydrogel precursor and NOA 73 pit layer cured with 0.5 $J/cm^2$ at 365nm (step 7 of Protocol D)</td></tr>
</table>

**Supplementary Table 2:** Curing conditions for the polymers and hydrogels.

### Supplementary Table 3:

| | **10x magnification** | **20x magnification** | **40x magnification** |
|---|---|---|---|
| **Size of a field view size** | 640 µm x1024 µm | 320 µm x 512 µm | 160 µm x 256 µm |
| **Min. feature size** | 2.3 µm | 1.2 µm | 0.6 µm |
| **Speed** | 4 min/mm2 | 6.1 min/mm2 | 16.6 min/mm2 |
| **Alignment precision** | Two patterns alignment depend on the motorized stage precision: **500 nm** | | |
| **Substrate** | Glass, Ibidi uncoated dish, NOA 73, MyPolymer 134 | | |
| **Protein incubation** | 5 min | | |
| **Protein type** | Fibrinogen, Laminin, Fibronectin, E-cadherin, Protein A, collagen I and IV (50 to 100 µg/ml). | | |

**Supplementary Table 3:** Characteristics of the illumination settings and size of the protein patterning of the micro-niches with PRIMO System.

#### Supplementary Note 1

### Fabrication of the pit layer membrane by soft lithography

The aim of the following protocol is to fabricate a negative PDMS (polydimethylsiloxane) mould of the pit. This PDMS mould will be further used as the final mould to fabricate the pit layer membrane in transparent biocompatible UV curable polymers (NOA 73 or MyPol-134).

**Fabrication of the PDMS pit mould**

1. Place the silicon wafer with the desired micro-well features in a vacuum jar with few drops (~2 - 400 µl) of Trichloro-perfluorooctyl-silane (Sigma, 448931) and expose the silicon wafer to the silane vapors for 4-6 hr.
2. Mix the PDMS elastomer base and the curing agent at a 10:1 ratio (w/w) and degas in a vacuum desiccator.
3. Pour the PDMS mixture over the master mould to achieve a 2 mm thick pit layer. Degas the PDMS layer before curing to let the PDMS mixture fill the microwells features of the wafer. Cure in the oven at 80 °C overnight.
4. Cut around the features to peel off the PDMS layer from the wafer.

**Fabrication of the pit layer membrane by soft lithography**

i. Cut a 5$mm^2$ square from the PDMS layer with inverse microwells features.
ii. Place the PDMS square onto a template flat surface (glass, plastic petri dish or PDMS) with the features side facing the surface and press at the 4 corners of the PDMS square to ensure full contact between the top of the PDMS features and the flat surface. This step is important to form a pit layer with through holes. **
iii. Place a drop of UV curable polymer on one of the four edge of the PDMS square and wait until the UV curable polymer fill the space between PDMS features by capillary effect (about 1 min for NOA 73 and 15 min for MyPolymer 134 for a 5 $mm^2$ PDMS square mould).
iv. Cure the UV curable polymer with the PDMS square under UV light (See conditions in **Supplementary Table 2**).
v. Peel off the PDMS mould from the cured polymer layer with tweezers. The fabricated pit layer membrane can be detached from the flat surface and used to assemble the desired micro-niches (See **Supplementary Table 1**).

** Critical step

#### Supplementary Note 2

### Protocols for Supplementary Table 1: Assembly and functionalization of the micro-niches

<u>**Protocol of Assembly of Pit and Base layers**</u>

**Assembly 1**

1. The pit layer is **directly** fabricated on the base layer.
2. Partially cured pit is required for *in situ* hydrogel growth. Please refer to Supplementary Table 1 for special curing conditions.

**Assembly 2**

1. The pit layer is fabricated on a template surface (glass, plastic petri dish or PDMS, See **Supplementary Note 1**).
2. Using tweezers, the pit layer is gently removed and transferred from the template onto the desired base surface.
3. If the base layer is fabricated with microstructures with different materials from the pit layer (e.g. NOA73 pit assemble with MYPolymer134 Base), air plasma treatment for the base and partially cured pit layers are needed to maintain good adhesiveness.
4. *Optional*: Seal two edges of the flattened membrane with a very tiny drop of NOA to create an edge, and cure it under UV light as previously.

**Assembly 3**

1. The pit layer is fabricated on a template surface (glass, plastic petri dish or PDMS).
2. The pit layer is coated with proteins on the template surface.
3. Cut around the microwells area but leave two opposite edges of polymer. This will help for the pit layer flipping.
4. The pit layer is gently removed, and importantly, **flipped before transferring** from the template onto the desired base surface. **
5. To ensure full contact between the microwells membrane and the coverslip, a flat PDMS layer can be used to press gently on the membrane.
6. The top surface of the pit layer, without any proteins, can then be passivated with antifouling agents (e.g. 0.2% Pluronic Acid diluted in PBS for 30 min).
7. *Optional*: Seal two edges of the flattened membrane with a very tiny drop of NOA 73 to create an edge, and cure it under UV light as previously.

<u>Notes</u>: To ensure a good adhesion between the polymer membrane and the coated plates, be sure that the plate is completely dry.

** Critical step

**Protocols for micro-niche functionalization**

**A – Homogeneous Protein Coating**

1. Substrates (glass, plastic, polymeric membranes or pillars), as pit or base layers, are subject to plasma treatment to promote physical adsorption of proteins.
2. Target proteins solutions (50µg/ml) were incubated at room temperature for 1h at RT (or 4˚C overnight). For pit layers, the protein solution is degased for 5min to avoid air bubbles trapped inside the microwells.
3. Remove protein solution and rinse with distilled water for 3 times.
4. Dry the dish with water or preserve with PBS.
5. Upon fully dried, the substrates are ready for assembly with another layer. Otherwise, the coated substrates can be stored in a freezer.

**B – Patterned Protein Coating**

1. Substrates (glass, plastic, polymeric membranes or pillars), as pit or base layers, are subject to oxygen plasma treatment to promote physical adsorption of PLL-g-PEG (PLL(20)-g[3.5]-PEG(5), 0.1 mg/ml in PBS pH=7.4, SuSoS AG, Dübendorf, Switzerland). **
2. After plasma treatment, the substrate is immediately incubated with a drop of 10 µL of PLL-g-PEG for at least 1 h at RT. For pit layers, degas the solution of PLL-g-PEG for 5min to avoid air bubbles trapped inside the wells. The solution can be hold with a PDMS sticker place around the pit membrane edge.
3. Rinse the substrate with PBS pH=7.4 for 5 times without drying the surface. **
4. Add 5 µL of photo-initiator solution PLPP (4-benzoylbenzyl-trimethylammonium chloride)
5. The target printing area is illuminated by DMD pattern-projected UV-laser (375 nm, PRIMO system, Alveole, France) for 40 s to cleave the PLL-g-PEG chain.
6. The substrate is rinsed with PBS pH=7.4 for 5 times.
7. Target proteins solutions in PBS (50µg/ml) are then incubated at room temperature for 5 min before rinsing with water for 3 times without drying the surface. **
8. If multiple proteins are needed, repeat steps 3 - 7 for the second proteins.
9. Dry the dish with water or preserve with PBS.

**C – (ECM) Hydrogels in situ Polymerization**

1. The pit layer made of partially cured NOA 73 is prepared as required (see curing conditions in **Supplementary Table 2** and fabrication **in Supplementary Note 1**).
2. A drop of (ECM) hydrogel precursor solution (better prepared on ice) is then directly added on top of the pits without any washing steps.
3. Degas for 5 min to allow the solution to enter the pit microwells and to avoid air bubbles trapped inside.
4. Load the substrate on to the stage of microscope with PRIMO system.
5. Open the live imaging function of the software (Metamorph) to observe and control the formation of hydrogels.
6. The target curing area, plus neighbouring regions that overlap the pit layers **, are illuminated by DMD pattern-projected UV-laser (375 nm, PRIMO). The curing conditions that depend on the types and concentrations of hydrogels are summarized in **Supplementary Table 2**. It is important to shine the pit area to induce the hydrogel *in situ* polymerization without photo-initiator.

7. The precursor solution is rinsed with PBS. Avoid extensive rinsing if the second type of hydrogels is to be patterned.
8. Fully cure the pit layers by UV light to avoid cytotoxicity.

Note: Cells could be encapsulated directly into the hydrogels by mixing precursor solutions with cell suspensions. In this case, the precursor solution is recommended to be prepared in CO2-independent media.
Note: For methacrylate Type I collagen gel, mix the precursor solution with 5% PEGDA. This will drastically improve the adhesion of hydrogels to pits.

**D – Homogeneous Hydrogel Base Protocol**

1. The base layer (glass coverslip) is modified with methacrylate groups by incubating 0.5% TMSPMA (3-(Trimethoxysilyl)propyl methacrylate) in ethanol diluted at 3% in 10% acetic acid in water (pH 3.5) for 5 min at RT. **
2. The substrate is rinsed with ethanol and dried.
3. To create a thin layer of hydrogels underneath, a 3 μL drop of hydrogel precursor solution without photo-initiator is added onto the TMSPMA treated glass coverslip.
4. The pit layer is fabricated on a template PDMS surface and partially cured by UV lamp (see curing conditions in **Supplementary Table 2**).
5. The partially-cured pit layer and the PDMS mould are carefully **removed together**** from the PDMS template surface and transferred onto the hydrogel precursors drop placed onto the TMSPMA treated glass coverslip (see step 1-3).
6. The hydrogel is cured with the pit layer by homogeneous UV illumination (see curing conditions in **Supplementary Table 2**). The hydrogel can therefore covalently bind to glass plate and the pit layer with strong adhesiveness.
7. The PDMS mould is gently peeled off from the pit layer with tweezers.

Note: If the protocol C for the pit is applied after the protocol D for the base, **special curing conditions** are applied. Indeed, a partially cured pit is required for in *situ* hydrogel growth and thus the curing time should be adjusted to allow fully curing of the hydrogel base but leave the pit layer partially cured. Please refer to **Supplementary Table 2** for this special curing condition.

** Critical step
